# Influenza A matrix protein M1 induces lipid membrane deformation via protein multimerization

**DOI:** 10.1101/565952

**Authors:** Ismail Dahmani, Kai Ludwig, Salvatore Chiantia

## Abstract

The matrix protein M1 of the Influenza A virus is supposed to mediate viral assembly and budding at the plasma membrane (PM) of infected cells. In order for a new viral particle to form, the PM lipid bilayer has to bend into a vesicle towards the extracellular side. Studies in cellular models have proposed that different viral proteins might be responsible for inducing membrane curvature in this context (including M1), but a clear consensus has not been reached. In this study, we use a combination of fluorescence microscopy, cryogenic transmission electron microscopy (cryo-TEM), cryo-electron tomography (cryo-ET) and scanning fluorescence correlation spectroscopy (sFCS) to investigate M1-induced membrane deformation in biophysical models of the PM. Our results indicate that M1 is indeed able to cause membrane curvature in lipid bilayers containing negatively-charged lipids, in the absence of other viral components. Furthermore, we prove that simple protein binding is not sufficient to induce membrane restructuring. Rather, it appears that stable M1-M1 interactions and multimer formation are required in order to alter the bilayer three-dimensional structure, through the formation of a protein scaffold. Finally, our results suggest that, in a physiological context, M1-induced membrane deformation might be modulated by the initial bilayer curvature and the lateral organization of membrane components (i.e. the presence of lipid domains).

## 1. Introduction

The influenza A virus (IAV) buds from the plasma membrane (PM) of infected cells and, while doing so, acquires a lipid envelope from its host (1–4). During this step, the PM lipid bilayer is initially bent into a vesicle, towards the extracellular side (5). Assembly of viral components, bending of the lipid bilayer and the resulting budding of virions are essential parts of the IAV replication cycle and, therefore, their regulation could be a potential target for antiviral therapies.

IAV assembly is thought to be orchestrated by its matrix protein M1, which mediates the interactions among other viral components and the lipids in the PM (6). In spite of its fundamental importance, this process is still not fully understood. We and others have investigated M1-M1 and M1-lipid interactions, both in model membranes and cellular systems (7–9). It was shown that M1 interacts electrostatically via its N-terminal domain with acidic lipids and this interaction modulates protein multimerization (7, 9, 10). On the other hand, the process by which M1 or other viral proteins can induce membrane curvature during IAV budding is less clear. This issue has been investigated *in cellulo* without reaching a clear consensus (11). In some cases, it was demonstrated that M1 alone is sufficient to induce budding at the PM, without the need for other viral components (12, 13). Other studies have shown instead that IAV hemagglutinin and neuraminidase are needed to produce viral particles, while M1 might not play a significant role (14–16). Another group suggested that, in the presence of significant M1-PM binding, the protein might be capable of inducing budding (17). In light of these observations, it would be therefore very interesting to investigate M1-lipid interactions in a controlled environment and clarify whether the protein has indeed the ability to induce curvature in lipid bilayers on its own. In this context, giant unilamellar vesicles (GUVs) provide a simplified model for the PM, consisting of a free-standing lipid bilayer with well-defined composition and physical properties (18–21). GUVs have been used in the past to investigate the interplay between viral proteins and lipid membranes and, specifically, protein-induced alterations in the three-dimensional structure of the bilayer (22–25).

In this work, we used an analogous approach to investigate the interaction between IAV M1 and lipid membranes, using a combination of fluorescence microscopy imaging, scanning fluorescence correlation spectroscopy (sFCS) (26), cryogenic transmission electron microscopy (cryo-TEM), as well as cryo-electron tomography (cryo-ET). Our results indicate that M1 is capable to cause membrane deformation, also in the absence of other proteins. Furthermore, we quantitatively demonstrate that M1 multimerization, rather than simple binding, is necessary for the induction of membrane curvature.

## 2. Materials and Methods

### 2.1 Chemicals

All lipids (i.e. dioleoyl-sn-glycero-3-phosphocholine (DOPC), 2-distearoyl-sn-glycero-3-phosphocholine (DSPC), 1,2-dioleoyl-sn-glycero-3-phospho-L-serine (DOPS), phosphatidylinositol 4,5-bisphosphate (PIP_2_), 1,2-dipalmitoyl-sn-glycero-3-phospho-(1′-rac-glycerol) (DPPG), cholesterol and 1,2-dioleoyl-*sn*-glycero-3-phosphoethanolamine-*N*-Lissamine rhodamine B sulfonyl (Rhodamine-DOPE)) were obtained from Avanti Polar Lipids (Alabaster, Alabama, USA). Alexa Fluor 488 succinimidyl ester was obtained from Life Technologies. (Darmstadt, Germany). 2-(4-(3-(4-acetyl-3-hydroxy-2-propylphenoxy) propoxy) phenoxy acetic acid (PHE) was purchased from Cayman Chemical (Ann Arbor, MI, USA). 10-fold concentrated phosphate buffer (PBS, 100 mM phosphate buffer, 27 mM potassium chloride and 1.37 M sodium chloride at pH 7.4) was from Alfa Aesar (Haverhill, MA, USA). Glucose was from VWR Chemicals (Radnor, PE, USA). All other chemicals were purchased from ROTH (Karlsruhe, Germany), unless differently specified. Indium tin oxide (ITO) coated cover slips, 20×20 mm, thickness #1, 8-12 Ohms resistivity, were purchased from SPI supplies (West Chester, PA, USA).

### 2.2 Preparation of giant unilamellar vesicles

Giant unilamellar vesicles (GUVs) were prepared using the electroformation method (27, 28). Different lipid compositions were used to prepare the vesicles, as specified in each case in the Results section. Typically, DOPC and cholesterol were mixed with various amounts of negatively charged lipids (e.g. DOPS) at molar ratios between 10 mol% and 50 mol%. 0.05 mol% or 0.1 mol% Rhodamine-DOPE was added as a tracer to allow GUV membrane visualization by confocal microscopy. The GUV electroformation chamber consisted of two conductive ITO coverslips facing each other and separated by a 3 mm thick teflon spacer. The total volume of the chamber was ~300 μL. 30 μL of a 3 mM lipid solution in chloroform and/or ethanol were spread on a preheated ITO coverslip, forming a thin lipid film. The solvent was evaporated using a nitrogen flow for 5 minutes at room temperature (note that solvent evaporation under vacuum for 1 h did not show a difference in vesicle behavior). After assembly, the chamber was filled with a sucrose solution in deionized water (e.g. 150mM) and connected to a voltage generator (AC generator FG 250 D, H-Tronic, Hirschau, Germany). For experiments at low pH, the chamber was filled with a 150 mM sucrose, 10 mM acetate buffer solution at pH 5. A sinusoidal electric field of 1.4 V at 10 Hz was applied for 1.5 h at room temperature. To facilitate the detachment of GUV from the slides, the voltage was decreased to 0.5 V for 30 minutes.

### 2.3 Preparation of GUVs showing phase separation

Vesicles were prepared as described in the previous paragraph, using a lipid mixture consisting of 10 mol% cholesterol, 45 mol% DOPC, 15 mol% DPPG and 30 mol% DSPC. Additionally, 0.05 mol% Rhodamine-DOPE was added to visualize the different lipid domains. Lipids were dissolved in chloroform/methanol 9:1 v:v (5 mM, prepared freshly and kept under a nitrogen atmosphere). The obtained lipid film was subjected to the electroformation procedure (see previous paragraph) at 50°C for 1.5 h. The chamber was then cooled down slowly at room temperature before imaging.

### 2.4 Expression and purification of recombinant matrix protein (M1) constructs

6xHis-tag M1 protein from Influenza A/FPV/Rostock/34 was expressed and purified, using a protocol adapted from Hilsch et al. (9). Rosetta *E. coli* (DE3) pLysS-competent cells were grown in 1 liter of medium (containing 0.2% glucose, 50 mg/mL chloramphenicol and 50 mg/mL ampicillin) until reaching OD_600_~0.7 at 37 °C. Then, protein expression was induced by addition of IPTG (0.4 mM) while shaking for 3 h at 37°C. Bacteria were harvested by centrifugation for 10 minutes at 4800 rpm (Thermo Lynx 4000 F12-6 rotor, Thermo Fisher Scientific, Waltham, MA, United States). The obtained pellets were stored at −80 °C until needed. On a different day, pellets were resuspended in lysis buffer (16 mM Na_2_HPO_4_, 146 mM KH_2_PO_4_, 500 mM NaCl, 5.4 mM KCl, EDTA-free protease inhibitor cocktail, 1 mM PMSF, 200 μg/ml DNAse), quickly frozen (15 minutes at −80°C) and incubated on a rotation shaker at 4°C for 30-60 minutes. All the following steps were performed at 4 °C. The bacteria were completely lysed using a French press (Glen Mills, NJ, USA) run at 1000 psi. Finally the obtained bacterial lysate was clarified by centrifugation at 21,000 rpm for 30 minutes (Thermo Lynx 4000, A27-8 rotor, Thermo Fisher Scientific, Waltham, MA, United States). The supernatant was incubated with 2 mL TALON resin (Takara, Saint-Germain-en-Laye, France) in a tube using a rotation shaker for 30-60 minutes. The resin was then washed with an equilibration buffer (8 mM Na_2_HPO_4_, 1.5 mM KH_2_PO_4_, 500 mM NaCl, 2.7 mM KCl, pH 7.4) and incubated on shaker for 15 minutes. The resin was washed again 2-3 times with an intermediate washing buffer consisting of the equilibration buffer with additional 60 mM imidazole, pH 7.2. Finally, M1 was eluted with an elution buffer (2-fold concentrated PBS with additional 180 mM imidazole, pH 7.4). Typical (maximal) protein concentrations were ~60 μM. SDS-PAGE was performed to verify protein purity. Protein concentration was determined using UV absorbance at 280 nm on an Agilent 8453 UV/Vis spectrophotometer (Agilent Technologies, Milford, MA, USA).

For the experiments performed at pH 5, the protein was purified directly at low pH. In particular, the intermediate washing buffer contained 20 mM imidazole (pH 6.5). The elution buffer contained 50 mM Sodium Acetate and 300 mM NaCl (pH 5).

M1-derived constructs (i.e. N-terminal domain 1-164 aa (NM1) and C-terminal domain 165-252 aa (CM1)) were purified at pH 7.4 as previously described (10). The elution buffer used for NM1 was the same as the one used for M1. The elution buffer used for CM1 was PBS with additional 150 mM imidazole (pH 7.4).

### 2.5 Protein labeling for fluorescence microscopy

The purified protein was conjugated, if needed, with the primary amine-reactive dye Alexa Fluor 488 succinimidyl ester. Freshly purified protein (in elution buffer) was incubated with the reactive dye (10:1 molar ratio) for 2 h at 4 °C, at the desired pH (7.4 or pH 5).

For simple imaging experiments, the protein-dye mixture was directly used, without further separation, as described in the next paragraph. For quantitative experiments (e.g. Par. 3.6), the labeled protein (M1-Alexa488) was separated from the free dye using size-exclusion chromatography (PD-10, GE Healthcare, Munich, Germany). The labeled protein was eluted with diluted PBS (pH 7.4), matched to the desired final osmolarity, as described in the next paragraph. Protein concentration and labeling efficiency were determined by absorbance at 280 nm and 490 nm, respectively. The maximum protein concentrations were typically around 40 μM, while labeling efficiencies ranged between 0.1 and 0.15 dye/protein.

### 2.6 Protein-GUV samples and imaging

Before mixing with GUVs, the protein solution (i.e. M1 in elution buffer) was diluted with deionized water to reach the target approximate osmolarity. For example, for experiments in which GUVs were prepared in 150 mM sucrose (Figs. 1, 3 and 4), the purified M1 solution (with or without Alexa Fluor 488 succinimidyl ester) was diluted ca. 5-fold. Typical protein concentrations at this step were ~12 μM. The protein solution was then mixed with the GUV suspension with different volume ratios, in order to obtain the final desired protein concentration (usually between 5 and 10 μM) in a total 300 μL volume, and transferred to 35-mm dishes (CellVis, Mountain View, CA) with optical glass bottoms, previously passivated with a 1% bovine serum albumin solution.

For the experiments carried in the presence of the M1-multimerization inhibitor PHE, M1 (~40 μM) was incubated with PHE (100 μM, unless differently stated) for 30 minutes directly in the elution buffer. Subsequently, the mixture was incubated with 4 μM Alexa Fluor 488 succinimidyl ester, for 2 hours at 4 °C. Finally, the protein solution was diluted with deionized water to approximatively match the osmolarity of the vesicle suspension and mixed with the GUVs, so to obtain a final protein concentration of 10 μM (and ~25 μM PHE).

Imaging experiments carried at pH 5 were carried out as follows: M1 was purified and labeled as described in Par. 2.4 and 2.5 directly at pH 5. The protein-dye solution in elution buffer (50 mM Sodium Acetate and 300 mM NaCl, pH 5) was subsequently diluted ~3 fold using deionized water, to a final protein concentration of ~13 μM (in ~16 mM sodium acetate, ~90 mM NaCl). This solution was finally mixed with the GUV suspension, so to obtain a final protein concentration of 10 μM.

We have verified that the presence of imidazole does not affect membrane curvature for at least 2 hours and that Alexa Fluor 488 succinimidyl ester does not significantly interact with GUVs (data not shown). Nevertheless, in the case of samples that required higher protein concentrations (i.e. >10 μM M1) or extensive removal of free dye, the elution buffer was exchanged after protein labeling, using a PD-10 column. For example, for the sFCS experiments described in Par. 3.6, the protein was eluted using 2-fold diluted PBS (pH 7.4), to approximatively match the osmolarity of GUV samples prepared in 150 mM sucrose.

Confocal fluorescence laser scanning microscopy (LSM) imaging was performed on a Zeiss LSM780 system (Carl Zeiss, Oberkochen, Germany) using a 40× 1.2 NA water-immersion objective. Samples were excited with a 488 nm argon laser (for the fluorophores Alexa Fluor 488 succinimidyl ester) or a 561 nm diode laser (for the fluorophore Rhodamine-DOPE). For measurements performed using 488 nm excitation, fluorescence was detected between 499 and 695 nm, after passing through a 488 nm dichroic mirror, using GaAsP detectors. For measurements performed with 561 nm excitation, fluorescence emission was separated using a 488/561 nm dichroic mirror and was detected between 570 and 695 nm.

### 2.7 Liposome preparation for cryo-transmission electron microscopy

Liposomes were prepared by extrusion. The desired lipids mixture (DOPC + 40 mol% DOPS) was dissolved in chloroform and then evaporated under nitrogen stream at room temperature to form a lipid film. Films were then rehydrated in PBS (pH 7.4). The obtained dispersion (2 mM total lipid concentration) was vigorously vortexed for 2–5 minutes. The diameter of the obtained multilamellar vesicles was reduced by serially extruding the suspension 12 times through a 100 nm pore diameter polycarbonate membrane (Whatman, Maidstone, UK) with a hand-held extruder (Avanti Polar Lipids, AL, USA). Liposome size (ca. 130 nm diameter in average) was measured via dynamics light scattering, using a Zetasizer HS 1000, (Malvern, Worcestershire, UK). The purified M1 (unlabeled) in elution buffer was diluted with deionized water to approximatively match the osmolarity of the liposome suspension (ca. 2.6-fold dilution). Liposomes and protein solution were mixed in a total volume of 800 μL so to obtain a 10 μM final M1 concentration. Typical final lipid concentrations were ~0.5-1 mM. Perforated carbon film-covered microscopical 200 mesh grids (R1/4 batch of Quantifoil, MicroTools GmbH, Jena, Germany) were cleaned with chloroform and hydrophilised by 60 s glow discharging at 8 W in a BAL-TEC MED 020 device (Leica Microsystems, Wetzlar, Germany), before 5 μl aliquots of the liposome/protein solution were applied to the grids. The samples were vitrified by automatic blotting and plunge freezing with a FEI Vitrobot Mark IV (Thermo Fisher Scientific, Waltham, USA) using liquid ethane as cryogen.

### 2.8 Cryo-transmission electron microscopy

The vitrified specimens were transferred under liquid nitrogen into the autoloader of a FEI TALOS ARCTICA electron microscope (Thermo Fisher Scientific, Waltham, USA). This microscope is equipped with a high-brightness field-emission gun (XFEG) operated at an acceleration voltage of 200 kV. Micrographs were acquired on a FEI Falcon 3 4k × 4k direct electron detector (Thermo Fisher Scientific, Waltham, USA) using a 70 μm objective aperture at a primary magnification of 28 k or 45 k, corresponding to a calibrated pixel size of 3.69 or 2.29 Å/pixel, respectively.

### 2.9 Cryo-electron tomography

Single axis tilt series (±60° in 2° tilt angle increments) were recorded with the Falcon 3 direct electron detector at full resolution (28 K primary magnification) with a total dose lower than 70 e-/Å^2^. Tomogram reconstruction was performed using ThermoFisher Inspect3D software. Amira, Version 6.0 (FEI, Oregon, USA) was used for visualization.

### 2.10 Scanning fluorescence correlation spectroscopy

Scanning fluorescence correlation spectroscopy (sFCS) measurements were performed on a Zeiss LSM780 system using a 40× 1.2 NA water-immersion objective. M1-Alexa488 was excited with a 488 nm argon laser. Fluorescence was detected after passing through a 520/35 nm bandpass filter. Data acquisition and analysis were performed as described in Dunsing et al. (26, 29). Briefly, line scans of 128 × 1 pixels (pixel size 160 nm) were performed perpendicular to the GUV membrane with a 472.73 μs scan time. Typically, 600,000 lines were acquired (total scan time ~4 minutes) in photon counting mode. Low laser power was used (~1 μW) to avoid photobleaching and fluorescence saturation effect. Data were exported as TIF files, imported and analyzed in Matlab (MathWorks, Natick, MA) using custom-written code (26, 29). The analysis of the thus obtained auto-correlation curves with a two-dimensional diffusion model resulted in the determination of the number of fluorophores in the confocal volume (N) and their apparent diffusion times. Furthermore, total fluorescence intensity at the GUV membrane and M1-Alexa488 brightness were determined. Protein brightness and fluorescence intensity were normalized to account for day-to-day variations in laser power, optics alignment and degree of protein labeling, as described in Hilsch et al. (9). An approximate conversion from normalized brightness to the size of protein multimers is discussed in Par. 3.6, assuming for the sake of simplicity that the lowest measured average brightness value (ca. 0.025 a.u., sample “pH 5”) corresponds to M1 dimers (30) and using the formula described in Dunsing et al. (31), with p_f_=0.1 and ɛ=0.0227.

Statistical significances of differences among sFCS data sets were determined using a two-sided t-test with distinct variances (ttest2 routine, Matlab).

## 3. Results

### 3.1 M1 induces membrane deformation in GUVs containing negatively charged lipids

In order to clarify whether M1 is sufficient to induce membrane deformation in protein-free lipid bilayers, we incubated GUVs of different compositions in the presence of 5 μM M1-Alexa448. It is worth noting that ca. 1 in 10 proteins were labeled (see Materials and Methods). Freshly-prepared lipid vesicles included increasing amounts of DOPS (from 0 to 50 mol%), as it was shown that this lipid promotes M1 binding via electrostatic interactions (7–10, 32). GUVs were stained with trace amounts of a fluorescent lipid (Rhodamine-DOPE) and observed via confocal LSM. According to Fig. 1 A-D and G, significant membrane deformation was observed for GUVs containing high amounts of DOPS (i.e. ≥30 mol%). GUVs containing low amounts of DOPS (e.g. 10 mol%, Fig. 1 B) or in control samples without protein (Fig. S1 A) remained mostly spherical. We did not observe significant changes in the shape of deformed GUVs during the measurement time.

Although alterations in membrane shape could be obtained also by using non-fluorescent M1 (Fig. S1 B), labeling of M1 was instrumental to directly visualize protein binding and organization. Figures 1 E-F and the inset in panel D show the spatial distribution of M1-Alexa488 typically observed in the samples represented in panels A, B and D, respectively. As expected, in the absence of DOPS, very little binding of M1 to the membrane was observed and most of the protein could be found in solution outside the GUVs (Fig. 1 E). In the case of GUVs containing 10 mol% DOPS, M1 bound homogeneously to the lipid membrane (Fig. 1 F). M1 binding appears therefore necessary but, in general, not sufficient to induce membrane deformation. Interestingly, we noticed that the protein bound homogeneously to non-spherical GUVs as well (e.g. Figure 1 D and inset). Also, the protein fluorescence intensity inside and outside deformed GUVs was, in general, not distinguishable (see e.g. inset in Fig. 1 D). This indicates that M1 might have crossed the membrane into the lumen of the vesicles in many cases. Nevertheless, we observed that the shape of deformed GUVs was qualitatively reproducible also whenever M1 was more clearly excluded from the lumen of the vesicles (see Fig. S1 B and C).

Additionally, an increase in membrane acyl-chain order (by increasing cholesterol concentration to 40 mol%, Fig. 1 H) did not appear to suppress membrane deformation. GUV shape alterations could also be observed if DOPS was substituted by other negatively-charged lipids, e.g. PIP_2_ or PG (data not shown). Finally, we observed that the N-terminal domain of M1 (M1N aa. 1-164) is sufficient to induce membrane deformation. The C-terminal domain (M1C aa. 165-252) shows no effect on GUV shape and, as previously reported (10), has a low degree of membrane binding (Fig. S2).

Table 1 (row “M1 pH 7.4”) shows a quantitative overview regarding the amounts of deformed GUVs observed for different compositions, in conditions similar to those relative to the samples shown in Fig. 1. In order to clearly detect protein binding and, thus, include in the quantification only larger GUVs (>~ 10μm diameter) that displayed a significant amount of bound M1, we increased the protein concentration to 10 μM (18 μM for GUVs with only 10 mol% DOPS). Visual inspection of several vesicles confirmed that membrane deformation is facilitated by higher DOPS concentrations. A quantitative characterization of M1 binding to deformed membranes is described below (Par 3.6)

**FIGURE 1:**
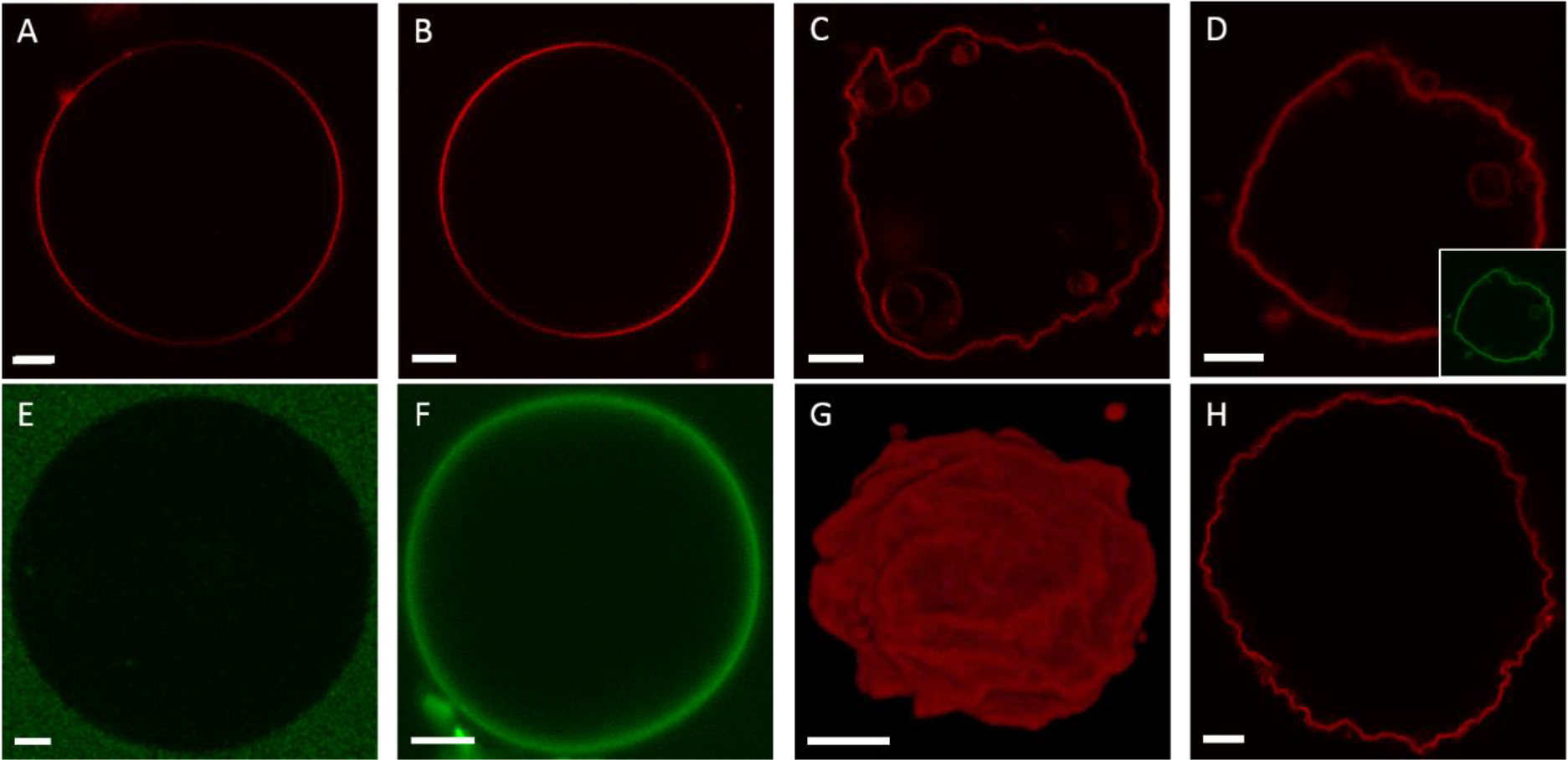
Shape alterations induced by M1 in DOPS-containing GUVs. **A-D**: Typical GUVs composed of 20 mol% Cholesterol, DOPC and increasing amounts of DOPS (A: 0 mol%, B: 10 mol%, C: 30 mol%, D: 50 mol%) observed via confocal LSM after 30 minutes incubation with 5 μM M1-Alexa488. Rhodamine-DOPE (0.01 mol%, red channel) was added to allow the visualization of the lipid bilayer via confocal LSM. **Inset in D, E-F**: Fluorescence signal originating from M1-Alexa488 (green channel) for typical GUVs containing 0 mol% DOPS (panel E, corresponding to the sample represented in panel A), 10 mol% DOPS (panel F, corresponding to the sample represented in panel B) and 50 mol% DOPS (inset in panel D). **G**: Three-dimensional reconstruction of a typical GUV containing 30 mol% DOPS in the presence of 5 μM M1-Alexa488 (corresponding to the sample represented in panel C). The fluorescence signal originates from Rhodamine-DOPE (0.01 mol%, red channel). **H**: Confocal LSM image of a typical GUV composed of 30 mol% DOPC, 30 mol% DOPS and 40 mol% cholesterol, in the presence of 5 μM M1-Alexa488. The fluorescence signal originates from Rhodamine-DOPE (0.01 mol%, red channel). All GUVs contained 150 mM sucrose in their lumen and were suspended in a phosphate buffered protein solution (pH 7.4) with similar osmolarity (see Materials and Methods). Scale bars are 5 μm. Images were acquired at 23°C.

### 3.2 M1-induced membrane bending in SUVs observed at high resolution

Characterization of bilayer deformation in GUVs via confocal LSM is limited by optical resolution. In order to obtain high-resolution information regarding the interplay between M1 binding and membrane curvature, we used cryo-TEM to observe SUVs in the presence of M1. Lipid vesicles containing DOPC and 40 mol% DOPS were incubated with 10 μM (unlabeled) M1 before freezing. The representative results shown in Fig. 2 A and B indicate that M1 binds to a large fraction of the lipid bilayers. By observing the apparent bilayer thickness, it is possible to distinguish regions of the bilayer with bound M1 (red and yellow arrows) from those devoid of protein (green arrows). In order to obtain a better insight into the conditions at the membrane, we have employed cryo-electron tomography (cryo-ET). Slices of the final 3D reconstruction, calculated from tilting series of images of the SUV embedded in vitreous ice, provide a more accurate representation of membrane spatial features, compared to individual cryo-TEM projection images. Fig. 2 C shows such a 15 nm thick section through the three-dimensional volume just in the middle of a SUV partially covered by the M1 protein. The protein-free bilayer has a thickness of about 4 nm. Bilayer regions which appear to be bound to M1 have a thickness between ca. 8 and 9 nm. Interestingly, the presence of M1 on vesicles is clearly correlated to changes in vesicle shape. Protein-free SUV surface regions display a positive mean curvature (referred to the membrane monolayer exposed to the protein solution) similar to that observed for control samples (Fig. S3). Membrane regions to which M1 has bound display various spatial features. Most often, we observed outward tubulation, i.e. membrane surfaces with zero Gaussian curvature, as indicated by the red arrows in Fig. 2. In other cases, we observed inward vesiculation or membrane regions with negative Gaussian curvature (yellow arrows in Fig. 2).

**FIGURE 2:**
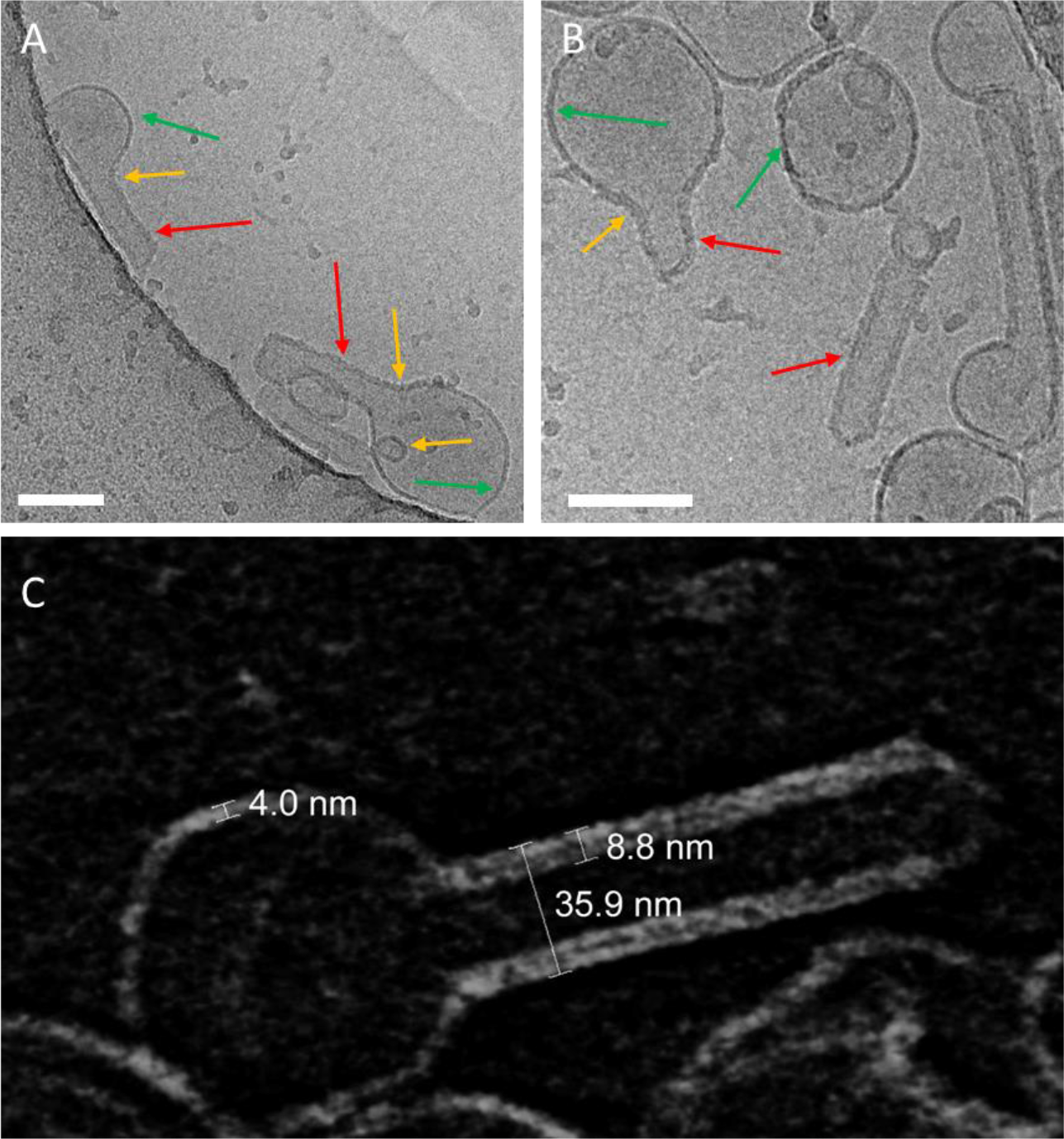
M1 modifies the curvature of SUV membranes. **A-B**: Typical cryo-TEM images of liposomes composed of 40 mol% DOPS in the presence of 10 μM M1 (see Materials and Methods). The arrows in panels A and B indicate protein-free membrane portions of vesicles (green arrows) or M1-bound portions of the bilayer which appear thicker (red arrows: tubules, yellow arrows tubule necks and inward vesiculation), due to the presence of bound M1. Scale bars are 100 nm. **C**: Cryo-electron tomography (cryo-ET) of a typical M1-bound liposome (tomography series ±60° at 2° angular increment): The image shows a 15 nm thick section through the three-dimensional volume just in the middle of a SUV incompletely bound to M1 (note the inverted contrast of the so-called voltex representation, i.e. lipid- and protein densities appear light). The numbers indicate the thickness of the bare lipid bilayer (ca. 4 nm), lipid bilayer with bound M1 (ca. 8-9 nm) and the diameter of a tubule originating from a vesicle (ca. 36 nm).

### 3.3 Lipid domains modulate M1-induced membrane bending

It was previously reported that IAV assembly and budding occurs in correspondence of confined PM domains (11). Furthermore, it is known that M1 binds to negatively charged lipids (7, 33). To investigate whether the spatial confinement of acidic lipids within membrane domains can modulate M1-induced membrane deformation, we produced GUVs displaying phase separation in ordered domains (i.e. bilayer regions characterized by highly-ordered lipid acyl chains, enriched in saturated phosphatidylcholine and saturated phosphatidylglycerol) and disordered domains (enriched in unsaturated phosphatidylcholine).

We have produced similar model lipid bilayers in the past by using a mixture of DOPC, DSPS, DSPC and cholesterol (34). When producing GUVs, we have noticed that exchanging the saturated phosphatidylserine with saturated phosphatidylglycerol (i.e., dipalmitoylphosphatidylglycerol, DPPG) improved the yield of phase-separated GUVs. Fig. 3 A shows an example of such GUVs observed via confocal microscopy (cholesterol:DPPG:DSPC:DOPC 10:15:30:45 molar ratios). The red channel refers to the lateral distribution of a fluorescent unsaturated lipid analogue (i.e. Rhodamine-DOPE), which strongly partitions into the disordered bilayer phase. The green channel refers to the distribution of a water-soluble fluorescent dye (Alexa Fluor 488 succinimidyl ester). The presence of the dye in the outer milieu allowed the visualization of the whole GUV shape. Ordered lipid domains can be thus simply identified by the low-partition of Rhodamine-DOPE and appear as dark membrane regions.

After addition of 10 μM M1-Alexa488, approximatively half the vesicles showed deviations from spherical shape. More in detail, we observed in certain cases M1-Alexa488 (green channel) bound to irregularly-shaped ordered domains (ca. 75% of the cases, typical domain size ~20±10 μm), as shown e.g. in Fig. 3 B. In other instances, we observed M1-Alexa488 bound to smaller ordered domains which were budding inwards (ca. 25% of the cases, typical domain size ~4±1 μm), as shown for example in Fig. 3 C. In agreement with previous observations (35, 36), we noticed that ordered domains sometime protruded outwards (see e.g. Fig. S4 A), independently of the presence of the protein. Finally, in the vast majority of cases, M1-Alexa488 appeared to be excluded from the lumen of the vesicles (see e.g. Fig. S4 B and C for enhanced contrast versions of Fig. 3 B and C).

**FIGURE 3:**
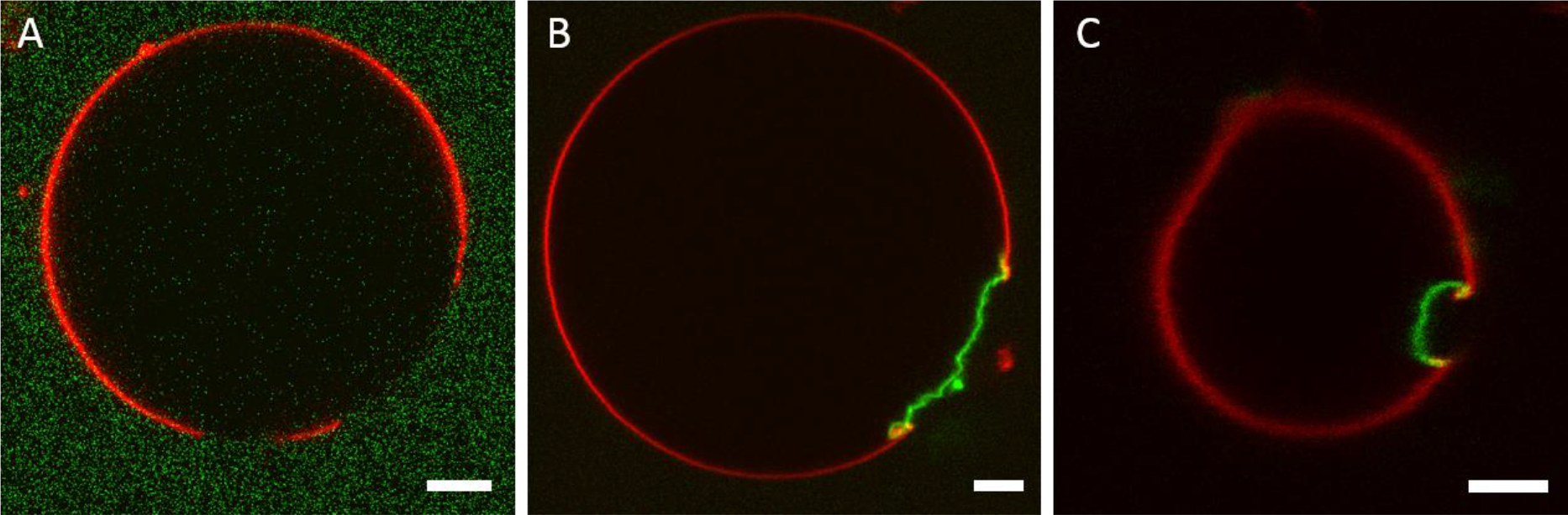
M1 binding to acidic lipid microdomains causes localized membrane deformation. **A**: Typical GUV composed of cholesterol:DPPG:DSPC:DOPC 10:15:30:45 molar ratios, imaged via confocal LSM. Phase separation was observed by labeling the vesicles with Rhodamine-DOPE 0.01 mol% (red channel, strongly enriched in the disordered phase) and introducing water-soluble Alexa Fluor 488 succinimidyl ester in the outer milieu of the vesicles (green channel). **B-C**: Typical GUVs with the same compositions as in panel A, in the presence of 10 μM M1-Alexa488. The protein is visualized in the green channel and the lateral distribution of Rhodamine-DOPE is represented in the red channel. GUVs similar to that shown in panel B were observed in ca. 75% of the cases. GUVs similar to that shown in panel C (inwards budding of the whole ordered domain) were observed in ca. 25% of the cases. All GUVs contained 150 mM sucrose in their lumen and were suspended in a phosphate buffered protein solution (pH 7.4) with similar osmolarity (see Materials and Methods). Scale bars are 5 μm. Images were acquired at 23°C.

### 3.4 Membrane deformation is accompanied by the formation of a stable M1-M1 network

So far, we have shown that M1-lipid interactions appear to causes alterations in the spatial organization of the bilayer. We next investigated whether M1-M1 interactions are also involved in the membrane deformation process. Fig. 4 A-B shows a typical GUV containing 30 mol% DOPS in the presence of 5 μM M1-Alexa488. As expected, within 30 minutes of incubation, deformed vesicles could be observed by monitoring the spatial distribution of a fluorescent lipid analogue (Fig. 4 B) or the labeled protein itself (Fig. 4 A). It is worth noting that the observed alterations in membrane shape are specific to M1. We have verified that extensive binding of another protein with high affinity to phosphatidylserine (i.e. Annexin V) does not cause significant membrane deformation (see Fig. S5).

We then proceeded to dissolve the GUVs, by adding 1.7 mM Triton X-100. Most of the lipids were effectively dissolved by the treatment, as demonstrated by the strong decrease of the fluorescence signal of the lipid analogue (Fig. 4 D, for an exemplar GUV). Interestingly, we observed that the protein was not affected by the detergent treatment, as it formed an apparently stable lipid-free three-dimensional structure (Fig. 4 C). We obtained similar results if lipid were dissolved using a mixture of different detergents (Triton X-100 1.7mM, n-Dodecylphosphocholine 1mM, CHAPS 10mM and n-Dodecyl-beta-Maltoside 1mM).

**FIGURE 4:**
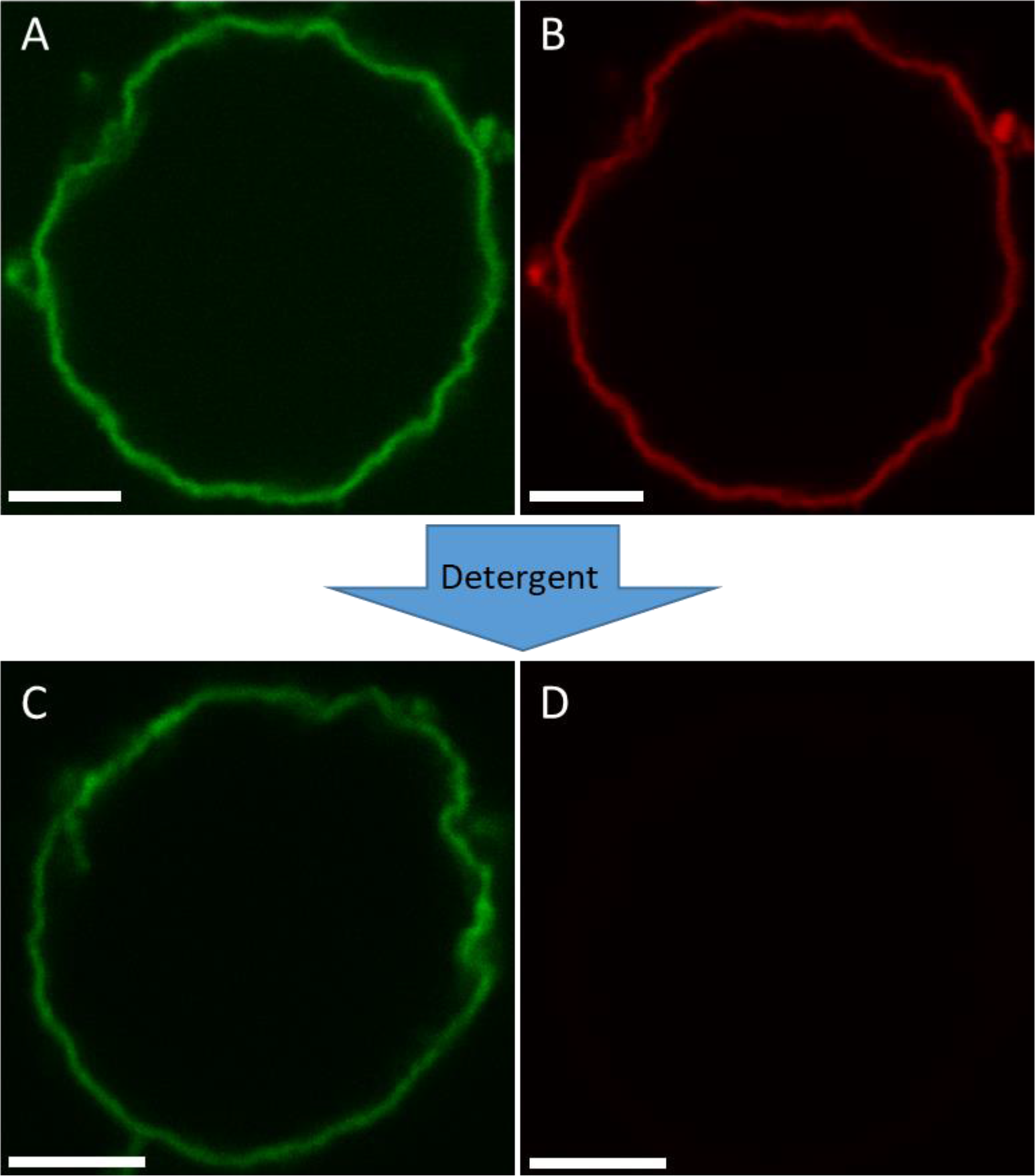
M1 layer interacting with the membrane is stable even after lipid removal. **A-B**: LSM confocal image of a typical GUV composed of DOPC:cholesterol:DOPS 50:20:30, after incubation with 5 μM M1-Alexa488 (green channel, panel A). The lipid bilayer is visualized via the addition of 0.05 mol% Rhodamine-DOPE (red channel, panel B). **C-D**: LSM confocal image of a (different) typical GUV after treatment for 5 minutes with detergent (e.g. Triton X-100 1.7 mM). Panel C represent the signal from M1-Alexa488. Panel D represents the signal from Rhodamine-DOPE. The excitation laser power used to acquire the image shown in panel D was ~7 times higher than the power used to acquire the image shown in panel B. All GUVs contained 150 mM sucrose in their lumen and were suspended in a phosphate buffered protein solution (pH 7.4) with similar osmolarity (see Materials and Methods). Scale bars are 5 μM. Images were acquired at 23°C.

### 3.5 M1-M1 interactions are needed for membrane deformation

The previous results suggest that M1 forms a protein network around GUVs that remains stable even after lipid removal. This observation hints at the presence of significant inter-protein interactions. To verify whether such M1-M1 interactions have a specific role in altering membrane curvature, we investigated conditions that inhibited protein multimerization, while not completely abolishing membrane binding. First, we incubated GUVs containing 40 mol% DOPS in the presence of 10 μM M1-Alexa488 pre-treated with 100 μM PHE. PHE is a small molecule that disrupts M1-M1 interactions, via direct interaction with the protein (37). The above-mentioned DOPS and M1 concentrations were chosen so that a strong deformation of the GUV bilayer would have been expected (compare to e.g. Fig. 1). Strikingly, most of the GUVs appeared spherical in the presence of PHE, as shown in Fig. 5 A. The same observation was made also under slightly different conditions (i.e. DOPS concentrations between 30 and 50 mol% and M1-Alexa488 incubated with 75-150 μM PHE, see Table 1).

Second, we incubated GUVs containing DOPC and 40 mol% DOPS in the presence of 10 μM M1-Alexa488 at pH 5. Low pH was already shown to interfere with M1 multimerization (9, 30, 38–42). Once again, we did not observe significant alterations in the shape of the GUVs (see e.g. Fig. 5 B). The same effect was observed for GUVs containing 30 mol% DOPS (see Table 1 for a quantitative summary).

**FIGURE 5:**
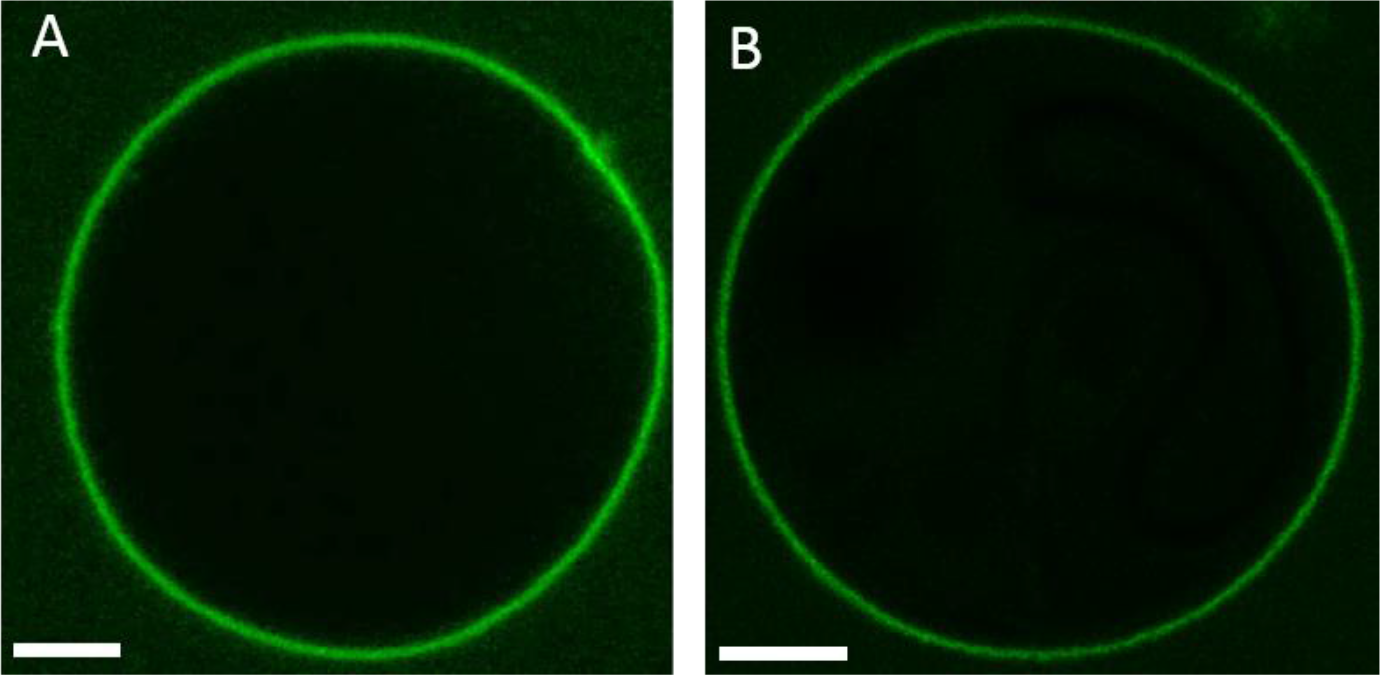
M1-induced membrane deformation requires M1 multimerization. **A**: Confocal LSM image of a typical GUV composed of 40 mol% DOPS and 60 mol% DOPC, in the presence of 10 μM fluorescent M1-Alexa488, which was pre-incubated with 100 μM PHE for 30 minutes. GUVs contained 150 mM sucrose in their lumen and were suspended in a phosphate buffered protein solution (pH 7.4) with similar osmolarity (see Materials and Methods). **B**: Confocal LSM image of a typical GUV with the same composition as in panel A, at pH 5, in the presence of 10 μM M1-Alexa488. These GUVs contained 150 mM sucrose, 10 mM sodium acetate buffer solution at pH 5 in their lumen. The external solution consisted of 10 μM M1-Alexa488 in ~15 mM sodium acetate buffer (pH 5) with ~70 mM NaCl buffer and ~30 mM sucrose (i.e. slightly hyperosmotic conditions, see Materials and Methods). Scale bars are 5 μm. Images were acquired at 23°C.

**Table 1:** Quantitative characterization of the amounts of non-spherical GUVs, for different membrane compositions. Percentages of deformed GUVs are reported for different experimental conditions discussed in the text. The row “M1 pH 7.4” refers to the conditions described for the experiments shown in Fig. 1. The row “M1 + PHE pH 7.4” refers to the experiments described in the context of Fig. 5 A. The observations reported in this table were performed by treating M1-Alexa488 with PHE concentrations between 75 and 150 μM. The row “M1 pH 5” refers to the experiments described in the context of Fig. 5 B. The total numbers of observed GUVs (n) refer in all cases to GUVs that were clearly labeled with fluorescent M1 and had a diameter >~ 10 μm. GUVs that did not display clearly recognizable M1-Alexa488 binding were not considered. The concentration of M1-Alexa488 was 10 μM, with the exceptions of the DOPS 10 mol% (18 μM) and DOPS 50 mol% (6 μM) samples. The percentages summarize the results of at least two independent experiments.

| **<u>GUV composition</u>** |  |  |  |  |  |
| --- | --- | --- | --- | --- | --- |
| DOPC + DOPS | DOPS 10 mol% | DOPS 20 mol% | DOPS 30 mol% | DOPS 40 mol% | DOPS 50 mol% |
|  | Relative amounts of deformed GUVs |  |  |  |  |
| **M1**<br>**pH 7.4** | **0%**<br>**(n=29)** | **24%**<br>**(n=33)** | **49%**<br>**(n=81)** | **83%**<br>**(n=84)** | **89%**<br>**(n=26)** |
| **M1 + PHE**<br>**pH 7.4** |  |  | **8%**<br>**(n=26)** | **13%**<br>**(n=23)** | **20%**<br>**(n=15)** |
| **M1**<br>**pH 5** | **0%**<br>**(n=46)** |  | **12%**<br>**(n=25)** | **17%**<br>**(n=23)** |  |

### 3.6 M1 exhibits reduced dynamics and high degree of multimerization in deformed vesicles

The results described in the previous paragraphs strongly suggest that M1 multimerization plays a determinant role in causing bilayer deformation. As an alternative approach to characterize protein-protein interactions, we performed scanning fluorescence correlation spectroscopy (sFCS) measurements on protein-bound GUVs. We have used in the past similar fluorescence fluctuation analysis methods to characterize M1 multimerization upon interaction with lipid bilayers (9). In this study, we used sFCS to directly quantify protein binding, dynamics and multimerization in spherical or deformed GUVs. It is worth noting that the sFCS approach (compared to e.g. point-FCS) is particularly suitable to investigate free-standing membranes. We incubated GUVs containing 30 mol% DOPS with 10 μM M1-Alexa488, in different conditions. First, we compared the properties of M1 bound to spherical or deformed GUV within a sample prepared at pH 7.4 (these measurements are referred to as “Spherical” and “Deformed”, respectively). As reported in Table 1, such samples contain in average ca. 50% deformed GUVs. Furthermore, we have performed sFCS on M1-Alexa488 bound to spherical GUVs in samples that were treated with the M1 multimerization inhibitor PHE (37) or that were prepared at pH 5 (these measurements are referred to as “PHE” and “pH 5”, respectively). Both conditions are supposed to be characterized by low M1 multimerization and reduced membrane deformation, as reported in the previous paragraph. Fig. 6 A shows, for the above-mentioned experimental conditions, the measured molecular brightness, i.e. a parameter indicative of to the clustering degree of the membrane-bound protein (9, 10). A normalization procedure was carried out to take into account day-to-day variations in protein labeling efficiency and in the experimental setup (see Par. 2.10). A conversion from molecular brightness to a precise multimeric state is, in this case, not straightforward. First, M1 is probably present as an undefined mixture of different multimeric species (and sFCS will detect a weighted average of the different multimeric species). Second, the low degree of protein labeling (ca. 0.1 label/protein ratio) implies that the effective number of fluorescent molecules in multimers of different sizes will be very similar. For example, fluorescent monomers and the vast majority of fluorescent dimers will both contain exactly one fluorescent label. This effect can be accounted for, as described by Dunsing et al. (31). Third, the brightness value of a monomeric reference would be required (i.e. a monomeric molecule labelled with the same fluorescent probe and diffusing in a system with the same geometrical properties as M1-Alexa488) (26). In this context, using the same fluorescent label diffusing in solution or bound to a different (monomeric) protein on the membrane would not be a precise reference in general, due to the different geometry (43) and possible changes in quantum yield of the probe, respectively. Nevertheless, some simplifications can be made in order to estimate the multimerization variations among the different samples. In the simple approximation that M1 is present i) as a single multimeric species and ii) in dimeric state at pH 5 (30), we could estimate that M1 forms approximatively decamers in spherical vesicles, 25-mers (und up to 100-mers) in deformed vesicles, and 15-mers in samples treated with PHE. These estimations take into account the degree of protein labeling, as described in Par. 2.10. In summary, our brightness measurements indicate that M1 bound to deformed vesicles (at pH 7.4, in the absence of PHE) is characterized by a significantly higher degree of multimerization, compared to the protein bound to spherical GUVs within the same samples. Also, both lower pH value and the presence of the multimerization inhibitor PHE resulted in significantly decreased M1 multimerization, as expected.

Additionally, sFCS provided quantitative information about M1-Alexa488 average diffusion dynamics. More precisely, we report the typical time needed by the protein to diffuse through a membrane area intersected by the detection volume. Such diffusion time is inversely proportional to the Brownian diffusion coefficient. The results shown in Fig. 6 B indicate, as expected, that M1 dynamics in deformed vesicles is slower than that observed in all other cases. It is worth noting that protein diffusion is, in general, dependent on both the size of protein assemblies and protein-membrane interaction.

In conclusion, these data indicate that M1-induced membrane deformation is clearly accompanied by a significant increase in protein multimerization and reduced protein dynamics. On the other hand, we did not observe a simple correlation between total amount of bound protein and the induction of membrane deformation, as the total fluorescence signal measured in deformed or spherical GUVs was similar (see Fig. S6).

**FIGURE 6:**
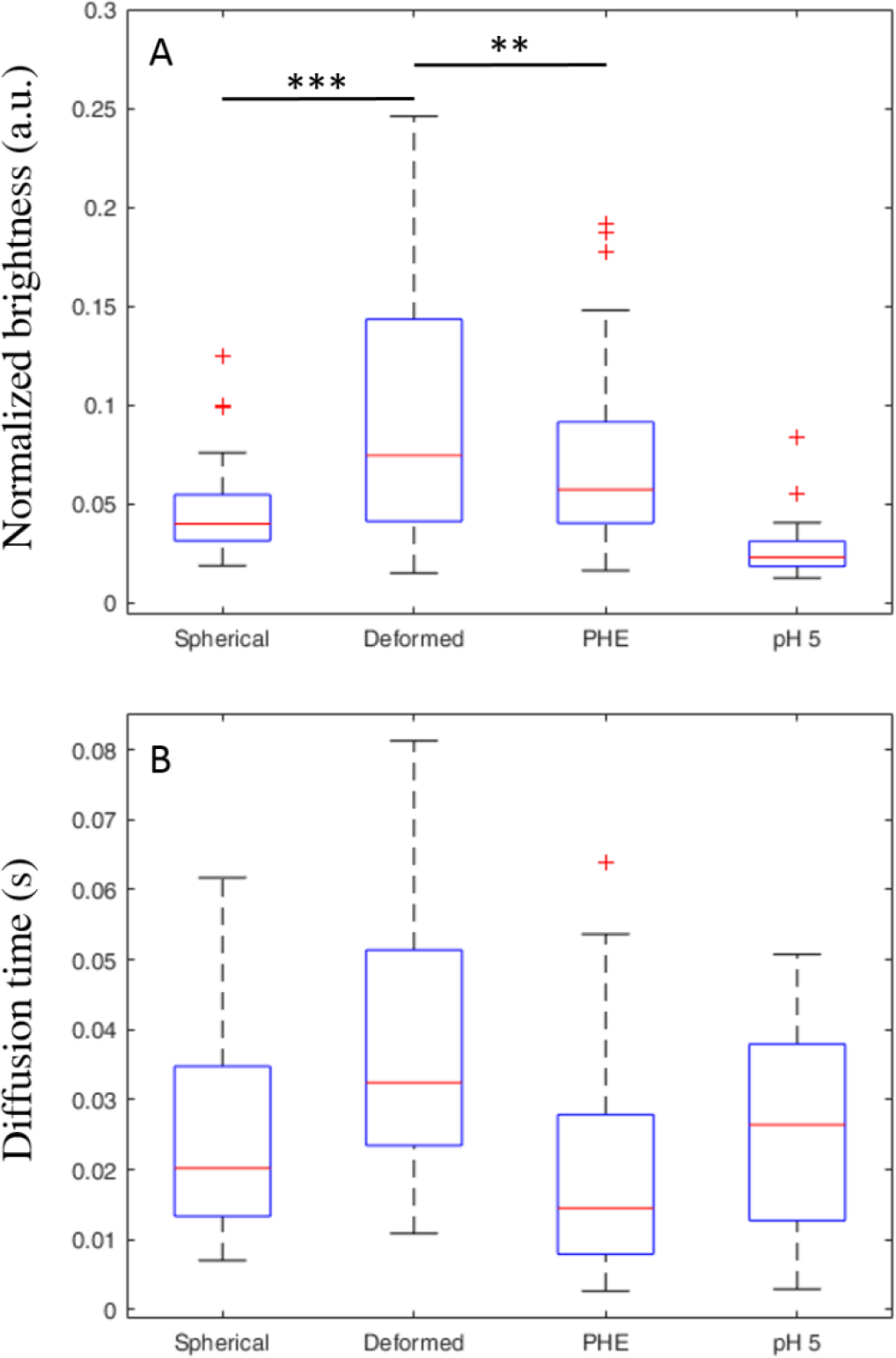
sFCS analysis of M1 brightness and dynamics in spherical and deformed GUVs. **A-B**: GUVs composed of 30 mol% DOPS and 70 mol% DOPC were incubated with 10 μM M1-Alexa488. The categories “Spherical” and “Deformed” refer to measurements in GUVs from samples at pH 7.4, in the absence of PHE. In these conditions, ca. 50% of the GUVs are clearly non-spherical (see Table 1). The category “PHE” refers to spherical GUVs in samples prepared at pH 7.4, using 100 μM PHE. In these conditions, ca. 90% of the GUVs are clearly spherical (see Table 1). The category “pH 5” refers to spherical GUVs in samples prepared at pH 5, in the absence of PHE. In these conditions, ca. 90% of the GUVs are clearly spherical (see Table 1). Details of sample preparation are described in the Materials and Methods section. sFCS measurements were performed on 16-34 GUVs from two independent sample preparations and the results were pooled together. Each measurement provided M1-Alexa488 normalized brightness values (shown as box plot in panel A), diffusion times (shown as box plot in panel B) and normalized fluorescence intensities (shown in Fig. S6). Upper outliers from the “Deformed” category are not included in the plot. “***” indicates statistical significant difference between categories, with a t-test probability outcome p<0.01. “**” indicates statistical significant difference between categories, with a t-test probability outcome p<0.05. In the case of diffusion time measurements (panel B), the category “Deformed” is significantly different from all the other categories, with a t-test probability outcome p<0.01, in all cases.

## 4. Discussion

The protein M1 is believed to play a fundamental role in the assembly of IAV (6). In order to release a new virion from the PM of an infected cell, the lipid bilayer has to undergo a shape change. In particular, some viral or cellular component(s) must induce a negative curvature on the inner leaflet of the PM, so that a virion can bud out from the cell. Previous studies have suggested that different viral proteins might be responsible for the reshaping of the PM (11–17). M1 has also been suggested to be capable of inducing curvature in the bilayer, but no unequivocal evidence has been presented yet. While investigations *in cellulo* provide information in a biologically relevant context (e.g. (12, 13)), the concurrent presence of several cellular and viral proteins does not allow the isolation and characterization of specific protein-protein or protein-lipid interactions. For this reason, we have applied here a *bottom up* approach and modelled the interaction between M1 and the PM in a controlled environment. In particular, we have characterized the interaction between M1 and physical models of the PM (i.e. GUVs or SUVs) using several microscopy methods. A similar approach has been recently used to investigate the interaction between the matrix protein from the Influenza C virus and lipid membranes (22). It was shown that this specific matrix protein forms elongated structures on lipid membranes and can also cause the formation of lipid tubules (by inducing negative membrane curvature). Nevertheless, the low sequence similarity between the two proteins and the morphologically different membrane structures observed in cells infected by the two viruses (44, 45) do not allow extending the conclusions regarding the matrix protein of Influenza C directly to Influenza A M1.

In the current work we have investigated for the first time the interaction between IAV M1 and free-standing model membranes containing different amounts of DOPS. The protein concentration (much larger than the reported K_d_ values (10, 39)) was chosen so to take into account that, *in vivo*, viral proteins might be recruited to small PM domains (46) and, therefore, reach a high local concentration (47). Previous investigations on solid-supported bilayers have demonstrated that M1 binds to lipid membranes containing negatively-charged lipids and that M1-lipid binding is accompanied by extensive protein multimerization (9). In agreement with these observations, we have observed that M1 binds effectively to GUVs containing DOPS. Surprisingly, M1 binding induced a significant alteration in the shape of the vesicles, especially at higher phosphatidylserine concentrations (e.g. >30 mol%, see Fig. 1). Specifically, the N-terminal domain of M1 is sufficient to induce membrane shape changes, in line with previous findings suggesting that this domain of the protein is responsible for M1-lipid and M1-M1 interactions (7, 10). In contrast, the N-terminal domain of Influenza C M1 is not sufficient to cause membrane curvature (22). Observation via optical microscopy indicates that M1 binds homogeneously while altering the spherical shape of vesicles. In general, both regions with positive and negative curvature can be observed. It is reasonable to assume that GUVs might become (temporarily) unstable, due to the significant and extensive deformation and volume decrease, thus allowing the internal solution to leak out. Accordingly, we have observed that M1 might indeed penetrate into the lumen of many vesicles and thus bind to both membrane leaflets. Nevertheless, reproducible membrane deformation patterns are observed also in the cases in which the protein seems to bind only to the outer leaflet (Fig. S1 or, in the case of spatially-limited deformations, Fig. 3 and S4). Alteration in bilayer curvature was also observed if M1 was incubated with SUVs with diameter ~100 nm. Thanks to the high spatial resolution of cryo-TEM and cryo-ET, M1-lipid interaction could be observed with remarkable detail. On a nanometer scale, binding of the protein to the vesicles appears not homogeneous. M1 seems capable to induce tubulation and, in fewer cases, vesiculation. The protein is found also concentrated in membrane regions with negative Gaussian curvature (i.e. at the neck of tubes) (Fig. 2). While comparing these results to the possible membrane-bending properties of M1 in a physiological context, it must be kept in mind that SUVs are highly curved compared to GUVs or the PM. The specific protein-lipid arrangements which are observed for SUVs (e.g. the ca. 20 nm radius observed for tubular structures) might therefore also be influenced by the initial membrane curvature. Furthermore, the protein-lipid ratio used in this study for GUV samples was at least 10 times lower than the one used for SUV sample, and this difference might play a role in the formation of protein-lipid structures. Nevertheless, taken together, the experiments performed in GUVs and SUVs suggest that M1 is sufficient to induce membrane deformation. While no other membrane component appears necessary for M1-induced membrane bending, the possibility that e.g. other viral proteins might modulate this effect (by, e.g., making negative curvature of the leaflet interacting with M1 energetically favorable over positive curvature) is currently under investigation. Furthermore, it is worth mentioning that M1 was reported to form various multimeric arrangements (e.g. helical (41)) that, *in vivo*, might result in a different remodeling of lipid membranes, compared to the one presented in the current work. The physical properties of the lipid bilayer do not appear to play a dominant role in M1-induced membrane bending, since we observed no effect induced by e.g. altering membrane fluidity (by increasing cholesterol content (48)), or due to changes in membrane lateral tension (due to varying osmotic pressure). Additionally, our observation that significant shape alteration is observed only in bilayers containing higher amounts of DOPS is likely not connected to an effect of this lipid on the physical properties of the membrane. It was suggested in fact that phosphatidylserine does not decrease the bending stiffness of a lipid bilayer (49, 50). In conclusion, the observed alterations in bilayer shape are brought about specifically by the binding of M1 to the membrane. Similar results are observed if M1 binds to GUVs via interactions with other lipids (i.e. in the absence of phosphatidylserine), such as PG, PIP_2_ or metal-ion-chelating lipids binding the His-tag of M1 (data not shown).

A further aspect that was examined in these experiments regarded the lateral organization of the bilayer, prior to protein binding. Previous investigations have shown that M1 multimerization and binding to lipid membranes are modulated by the presence of lipid domains containing negatively-charged lipids (34). In infected cells, M1 clusters are observed in correspondence of phosphatidylserine-rich membrane regions (34). In order to clarify whether M1-induced membrane restructuring is also affected by the lateral organization of the lipid bilayer, we investigated lipid mixtures displaying phase separation. In particular, GUVs contained ordered domains (enriched in cholesterol, saturated lipids and, reasonably, negatively-charged lipids) in a disordered bilayer (enriched in unsaturated lipids). In the presence of M1, the ordered domains showed variations from the original spherical shape and, especially in the case of smaller domains, inward budding (see e.g. Fig. 3 B-C). This observation demonstrates, on one hand, that these ordered domains are indeed enriched in acidic lipids to which M1 can effectively bind. Second, it indicates that the membrane restructuring effect of M1 appears significant even at relatively high values of membrane bending rigidity (cfr., for example, ca. 65 kT for a DPPC:cholesterol 80:20 bilayer (51)). Third, it suggests that the local membrane restructuring induced by M1 might be modulated by the lateral lipid organization in the bilayer region in which M1 is concentrated. This phenomenon might be relevant *in vivo*, since M1 is supposed to be confined in budozones, i.e. small domains of the PM of infected cells from which IAV budding takes place (52).

Finally, we aimed to clarify the molecular mechanism driving the deformation of the membrane induced by M1. Several mechanisms for the induction of membrane curvature driven by (non trans-membrane) proteins were described, including: membrane insertion, protein crowding and scaffolding (53–56). In order to distinguish among these possibilities, we observed M1-GUV interaction in conditions in which M1 could bind to the membrane but protein multimerization was hindered (i.e. low pH or in the presence of a multimerization inhibitor, see Fig. 5). These experiments revealed that protein binding to the bilayer *per se* did not induce significant alterations in membrane shape. The reduced ability of M1 to form large multimers in these conditions was confirmed via sFCS measurements (Fig. 6). Accordingly, the amount of M1 bound to deformed vesicles was not significantly higher than that of M1 bound to spherical vesicles (Fig. S6). Finally, we verified that binding of another protein to DOPS in GUVs (in comparable amounts to M1) did not affect membrane shape (Fig. S5). Taken together, these observations suggest that membrane insertion or protein crowding are not the main factors driving M1-induced curvature. On the other hand, we observed that M1 forms a protein network which is characterized by slow dynamics (Fig. S5, Fig. 6 B) and seems to impose its irregular (corrugated) shape on the underlying bilayer. The stability of M1-M1 interactions allows the protein shell to remain stable even after removal of the lipid membrane using detergents (Fig. 4). Additionally, we were able to quantitatively verify that M1 bound to deformed vesicles forms in general larger multimers (up to ca. 10-fold), compared to the case of spherical vesicles (Fig. 6). At this regard, the limitations of the sFCS approach in this context should be mentioned: First, the reported brightness/multimerization and diffusion values refer to an average of different multimeric species that might be present in the sample. Second, the presence of an immobile protein fraction would not be detected by fluorescence fluctuations techniques which, in general, report only of the properties of diffusing molecules. Third, membrane geometry and the detection area are not well-defined in the case of deformed membranes with large curvature (compared to the typical size of the detection volume of ca. 300 nm). In other words, a larger than expected bilayer surface (due to ruffling within the detection volume) might be observed during our experiments. As a consequence, protein diffusion times and total fluorescence intensity might be overestimated in deformed vesicles. On the other hand, protein brightness and multimerization would be underestimated. Of interest, these limitations do not affect our main findings that: i) the amount of M1 bound to deformed vesicles is not significantly higher than that of M1 bound to spherical vesicles and ii) M1 bound to deformed vesicles is characterized by a higher degree of multimerization.

In conclusion, our data show that IAV M1 binds to vesicles containing negatively-charged lipids, forming a stable protein layer. This M1-M1 network seems to be characterized by both regions of negative and positive curvature, although it appears that its precise spatial structure might be modulated by the initial curvature of the bilayer and the presence of lipid-protein lateral confinement. Our results support the model according to which the M1 shell, *in vivo*, might drive the IAV budding (likely in concert with other membrane and/or viral components that help determining a specific curvature) and provides mechanical stability to the newly formed virion (see e.g. (57)).

## Acknowledgments

The authors thank Rumiana Dimova for reading the manuscript and providing useful suggestions. This work was supported by the German Research Foundation (DFG grant CH 1238/3 to SC).

## References

1. Rossman JS, Lamb RA. 2011. Influenza virus assembly and budding. Virology 411:229–236.

2. Dou D, Revol R, Ostbye H, Wang H, Daniels R. 2018. Influenza A Virus Cell Entry, Replication, Virion Assembly and Movement. Front Immunol 9:1581.

3. Nayak DP, Hui EK, Barman S. 2004. Assembly and budding of influenza virus. Virus Res 106:147–165.

4. Nayak DP, Balogun RA, Yamada H, Zhou ZH, Barman S. 2009. Influenza virus morphogenesis and budding. Virus Res 143:147–161.

5. McMahon HT, Gallop JL. 2005. Membrane curvature and mechanisms of dynamic cell membrane remodelling. Nature 438:590–596.

6. Kordyukova LV, Shtykova EV, Baratova LA, Svergun DI, Batishchev OV. 2018. Matrix proteins of enveloped viruses: a case study of Influenza A virus M1 protein. J Biomol Struct Dyn doi:10.1080/07391102.2018.1436089:1–20.

7. Baudin F, Petit I, Weissenhorn W, Ruigrok RW. 2001. In vitro dissection of the membrane and RNP binding activities of influenza virus M1 protein. Virology 281:102–108.

8. Ruigrok RW, Barge A, Durrer P, Brunner J, Ma K, Whittaker GR. 2000. Membrane interaction of influenza virus M1 protein. Virology 267:289–298.

9. Hilsch M, Goldenbogen B, Sieben C, Hofer CT, Rabe JP, Klipp E, Herrmann A, Chiantia S. 2014. Influenza A matrix protein M1 multimerizes upon binding to lipid membranes. Biophys J 107:912–923.

10. Hofer CT, Di Lella S, Dahmani I, Jungnick N, Bordag N, Bobone S, Huang Q, Keller S, Herrmann A, Chiantia S. 2019. Structural determinants of the interaction between influenza A virus matrix protein M1 and lipid membranes. Biochim Biophys Acta Biomembr 1861:1123–1134.

11. Veit M, Thaa B. 2011. Association of influenza virus proteins with membrane rafts. Adv Virol 2011:370606.

12. Gomez-Puertas P, Albo C, Perez-Pastrana E, Vivo A, Portela A. 2000. Influenza virus matrix protein is the major driving force in virus budding. J Virol 74:11538–11547.

13. Latham T, Galarza JM. 2001. Formation of wild-type and chimeric influenza virus-like particles following simultaneous expression of only four structural proteins. J Virol 75:6154–6165.

14. Chen BJ, Leser GP, Morita E, Lamb RA. 2007. Influenza virus hemagglutinin and neuraminidase, but not the matrix protein, are required for assembly and budding of plasmid-derived virus-like particles. J Virol 81:7111–7123.

15. Chlanda P, Schraidt O, Kummer S, Riches J, Oberwinkler H, Prinz S, Krausslich HG, Briggs JA. 2015. Structural Analysis of the Roles of Influenza A Virus Membrane-Associated Proteins in Assembly and Morphology. J Virol 89:8957–8966.

16. Lai JC, Chan WW, Kien F, Nicholls JM, Peiris JS, Garcia JM. 2010. Formation of virus-like particles from human cell lines exclusively expressing influenza neuraminidase. J Gen Virol 91:2322–2330.

17. Wang D, Harmon A, Jin J, Francis DH, Christopher-Hennings J, Nelson E, Montelaro RC, Li F. 2010. The lack of an inherent membrane targeting signal is responsible for the failure of the matrix (M1) protein of influenza A virus to bud into virus-like particles. J Virol 84:4673–4681.

18. Walde P, Cosentino K, Engel H, Stano P. 2010. Giant vesicles: preparations and applications. Chembiochem 11:848–865.

19. Sezgin E, Schwille P. 2012. Model membrane platforms to study protein-membrane interactions. Mol Membr Biol 29:144–154.

20. Lagny TJ, Bassereau P. 2015. Bioinspired membrane-based systems for a physical approach of cell organization and dynamics: usefulness and limitations. Interface Focus 5:20150038.

21. Dimova R, Aranda S, Bezlyepkina N, Nikolov V, Riske KA, Lipowsky R. 2006. A practical guide to giant vesicles. Probing the membrane nanoregime via optical microscopy. J Phys Condens Matter 18:S1151–1176.

22. Saletti D, Radzimanowski J, Effantin G, Midtvedt D, Mangenot S, Weissenhorn W, Bassereau P, Bally M. 2017. The Matrix protein M1 from influenza C virus induces tubular membrane invaginations in an in vitro cell membrane model. Sci Rep 7:40801.

23. Solon J, Gareil O, Bassereau P, Gaudin Y. 2005. Membrane deformations induced by the matrix protein of vesicular stomatitis virus in a minimal system. J Gen Virol 86:3357–3363.

24. Soni SP, Stahelin RV. 2014. The Ebola virus matrix protein VP40 selectively induces vesiculation from phosphatidylserine-enriched membranes. J Biol Chem 289:33590–33597.

25. Shnyrova AV, Ayllon J, Mikhalyov II, Villar E, Zimmerberg J, Frolov VA. 2007. Vesicle formation by self-assembly of membrane-bound matrix proteins into a fluidlike budding domain. J Cell Biol 179:627–633.

26. Dunsing V, Chiantia S. 2018. A Fluorescence Fluctuation Spectroscopy Assay of Protein-Protein Interactions at Cell-Cell Contacts. J Vis Exp doi:10.3791/58582.

27. Angelova MI, Dimitrov DS. 1986. Liposome Electroformation. Faraday Discussions 81:303–+.

28. Mauroy C, Portet T, Winterhalder M, Bellard E, Blache MC, Teissie J, Zumbusch A, Rols MP. 2012. Giant lipid vesicles under electric field pulses assessed by non invasive imaging. Bioelectrochemistry 87:253–259.

29. Dunsing V, Mayer M, Liebsch F, Multhaup G, Chiantia S. 2017. Direct evidence of amyloid precursor-like protein 1 trans interactions in cell-cell adhesion platforms investigated via fluorescence fluctuation spectroscopy. Mol Biol Cell 28:3609–3620.

30. Zhang K, Wang Z, Liu X, Yin C, Basit Z, Xia B, Liu W. 2012. Dissection of influenza A virus M1 protein: pH-dependent oligomerization of N-terminal domain and dimerization of C-terminal domain. PLoS One 7:e37786.

31. Dunsing V, Luckner M, Zuhlke B, Petazzi RA, Herrmann A, Chiantia S. 2018. Optimal fluorescent protein tags for quantifying protein oligomerization in living cells. Sci Rep 8:10634.

32. Das SC, Watanabe S, Hatta M, Noda T, Neumann G, Ozawa M, Kawaoka Y. 2012. The highly conserved arginine residues at positions 76 through 78 of influenza A virus matrix protein M1 play an important role in viral replication by affecting the intracellular localization of M1. J Virol 86:1522–1530.

33. Kerviel A, Dash S, Moncorge O, Panthu B, Prchal J, Decimo D, Ohlmann T, Lina B, Favard C, Decroly E, Ottmann M, Roingeard P, Muriaux D. 2016. Involvement of an Arginine Triplet in M1 Matrix Protein Interaction with Membranes and in M1 Recruitment into Virus-Like Particles of the Influenza A(H1N1)pdm09 Virus. PLoS One 11:e0165421.

34. Bobone S, Hilsch M, Storm J, Dunsing V, Herrmann A, Chiantia S. 2017. Phosphatidylserine Lateral Organization Influences the Interaction of Influenza Virus Matrix Protein 1 with Lipid Membranes. J Virol 91.

35. Bacia K, Schwille P, Kurzchalia T. 2005. Sterol structure determines the separation of phases and the curvature of the liquid-ordered phase in model membranes. Proc Natl Acad Sci U S A 102:3272–3277.

36. Baumgart T, Hess ST, Webb WW. 2003. Imaging coexisting fluid domains in biomembrane models coupling curvature and line tension. Nature 425:821–824.

37. Mosier PD, Chiang MJ, Lin Z, Gao Y, Althufairi B, Zhou Q, Musayev F, Safo MK, Xie H, Desai UR. 2016. Broad Spectrum Anti-Influenza Agents by Inhibiting Self-Association of Matrix Protein 1. Sci Rep 6:32340.

38. Zhirnov OP. 1992. Isolation of matrix protein M1 from influenza viruses by acid-dependent extraction with nonionic detergent. Virology 186:324–330.

39. Batishchev OV, Shilova LA, Kachala MV, Tashkin VY, Sokolov VS, Fedorova NV, Baratova LA, Knyazev DG, Zimmerberg J, Chizmadzhev YA. 2016. pH-Dependent Formation and Disintegration of the Influenza A Virus Protein Scaffold To Provide Tension for Membrane Fusion. J Virol 90:575–585.

40. Zhirnov OP. 1990. Solubilization of matrix protein M1/M from virions occurs at different pH for orthomyxo- and paramyxoviruses. Virology 176:274–279.

41. Shtykova EV, Baratova LA, Fedorova NV, Radyukhin VA, Ksenofontov AL, Volkov VV, Shishkov AV, Dolgov AA, Shilova LA, Batishchev OV, Jeffries CM, Svergun DI. 2013. Structural analysis of influenza A virus matrix protein M1 and its self-assemblies at low pH. PLoS One 8:e82431.

42. Shtykova EV, Dadinova LA, Fedorova NV, Golanikov AE, Bogacheva EN, Ksenofontov AL, Baratova LA, Shilova LA, Tashkin VY, Galimzyanov TR, Jeffries CM, Svergun DI, Batishchev OV. 2017. Influenza virus Matrix Protein M1 preserves its conformation with pH, changing multimerization state at the priming stage due to electrostatics. Sci Rep 7:16793.

43. Macdonald PJ, Chen Y, Wang X, Chen Y, Mueller JD. 2010. Brightness analysis by Z-scan fluorescence fluctuation spectroscopy for the study of protein interactions within living cells. Biophys J 99:979–988.

44. Nishimura H, Hara M, Sugawara K, Kitame F, Takiguchi K, Umetsu Y, Tonosaki A, Nakamura K. 1990. Characterization of the cord-like structures emerging from the surface of influenza C virus-infected cells. Virology 179:179–188.

45. Muraki Y, Washioka H, Sugawara K, Matsuzaki Y, Takashita E, Hongo S. 2004. Identification of an amino acid residue on influenza C virus M1 protein responsible for formation of the cord-like structures of the virus. J Gen Virol 85:1885–1893.

46. Leser GP, Lamb RA. 2017. Lateral Organization of Influenza Virus Proteins in the Budozone Region of the Plasma Membrane. J Virol 91.

47. McLaughlin S, Wang J, Gambhir A, Murray D. 2002. PIP(2) and proteins: interactions, organization, and information flow. Annu Rev Biophys Biomol Struct 31:151–175.

48. Henriksen J, Rowat AC, Ipsen JH. 2004. Vesicle fluctuation analysis of the effects of sterols on membrane bending rigidity. Eur Biophys J 33:732–741.

49. Song J, Waugh RE. 1990. Bilayer membrane bending stiffness by tether formation from mixed PC-PS lipid vesicles. J Biomech Eng 112:235–240.

50. Dimova R. 2014. Recent developments in the field of bending rigidity measurements on membranes. Adv Colloid Interface Sci 208:225–234.

51. Doktorova M, Harries D, Khelashvili G. 2017. Determination of bending rigidity and tilt modulus of lipid membranes from real-space fluctuation analysis of molecular dynamics simulations. Phys Chem Chem Phys 19:16806–16818.

52. Schmitt AP, Lamb RA. 2005. Influenza virus assembly and budding at the viral budozone. Adv Virus Res 64:383–416.

53. McMahon HT, Boucrot E. 2015. Membrane curvature at a glance. J Cell Sci 128:1065–1070.

54. Bassereau P, Jin R, Baumgart T, Deserno M, Dimova R, Frolov VA, Bashkirov PV, Grubmuller H, Jahn R, Risselada HJ, Johannes L, Kozlov MM, Lipowsky R, Pucadyil TJ, Zeno WF, Stachowiak JC, Stamou D, Breuer A, Lauritsen L, Simon C, Sykes C, Voth GA, Weikl TR. 2018. The 2018 biomembrane curvature and remodeling roadmap. J Phys D Appl Phys 51.

55. Zimmerberg J, Kozlov MM. 2006. How proteins produce cellular membrane curvature. Nat Rev Mol Cell Biol 7:9–19.

56. Stachowiak JC, Hayden CC, Sasaki DY. 2010. Steric confinement of proteins on lipid membranes can drive curvature and tubulation. Proc Natl Acad Sci U S A 107:7781–7786.

57. Schaap IA, Eghiaian F, des Georges A, Veigel C. 2012. Effect of envelope proteins on the mechanical properties of influenza virus. J Biol Chem 287:41078–41088.

