## Supplementary Info for "Influenza A matrix protein M1 induces lipid membrane deformation via protein multimerization"

### SUPPLEMENTARY MATERIAL to Dahmani et al.

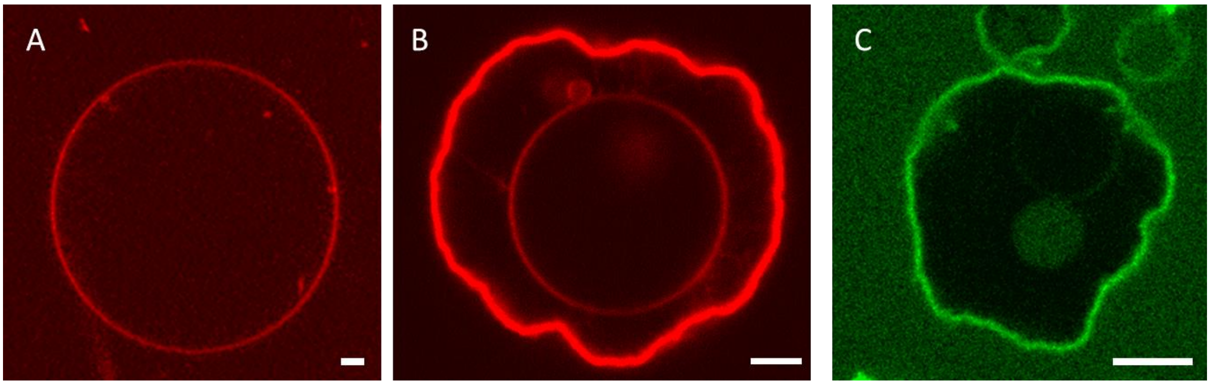


**FIGURE S1: Control experiments regarding the effect of labeling and buffer conditions.**

**A**: Confocal LSM image of a typical GUV composed of DOPC:DOPS 70:30 molar ratios. These GUVs contained 150 mM sucrose in their lumen and were suspended in a phosphate buffered solution pH 7.4 with similar osmolarity (exactly as for the samples shown in Fig.1 of the main text), but without M1. The lipid bilayer was labeled with 0.01 mol% Rhodamine-DOPE (red channel). In these conditions, all the GUVs we have examined (n=41) appeared spherical in shape. The conclusion that M1 (rather than e.g. buffer conditions) is uniquely responsible for membrane deformation can also be indirectly deduced from other results, such as those shown in Fig. S2 A or Fig. 5.

**B**: Confocal LSM image of a GUV composed of DOPC:DOPS 50:50 molar ratios, in the presence of 10 µM unlabeled M1. The lipid bilayer was labeled with 0.01 mol% Rhodamine-DOPE (red channel). GUVs contained 150 mM sucrose in their lumen and were suspended in a phosphate buffered protein solution (pH 7.4) with similar osmolarity (see Materials and Methods). This specific (deformed) vesicle contains a second GUV in its lumen. Such concentric vesicles could be observed in several instances. The fact that the internal GUV remained spherical indirectly suggests that this vesicle did not come in contact with a significant amount of protein. In other words, it is reasonable to assume that, in this case, M1 did not enter in the lumen of the deformed GUV.

**C**: Confocal LSM image of a GUV composed of DOPC:DOPS 70:30 molar ratios, in the presence of 2 µM M1-Alexa488 (green channel). GUVs contained 150 mM sucrose in their lumen and were suspended in a phosphate buffered protein solution (pH 7.4) with similar osmolarity (see Materials and Methods). In this specific case (representative of ca. 10% of the observed GUVs), it is possible to observe that the concentration of M1-Alexa488 is very low in the lumen. In other words, the protein is probably bound only to the outer leaflet of the GUV. Nevertheless, the general shape of the deformed vesicle is comparable to other cases (see e.g. Fig. 1 of the main text) for which the presence of M1 in the GUV lumen cannot be excluded.

Scale bars are 5 µm. Images were acquired at 23°C.

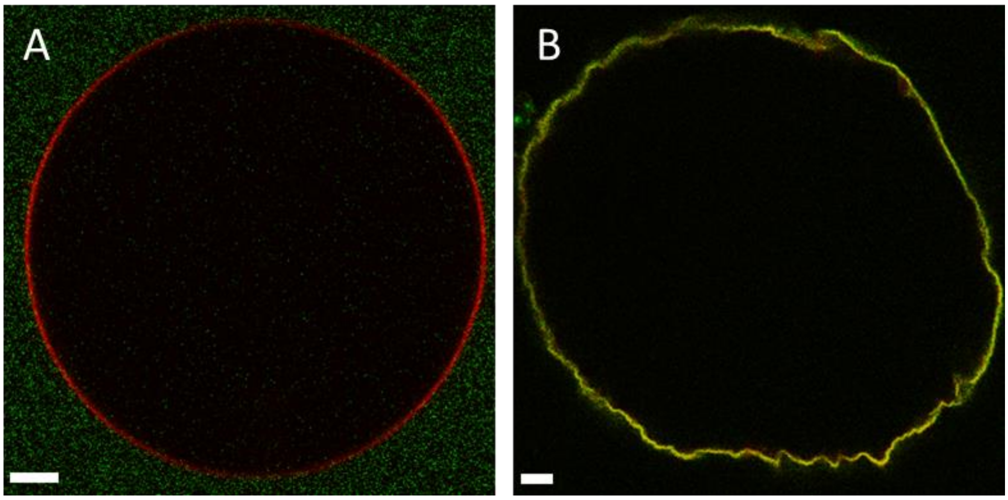


**FIGURE S2: The N-terminal domain of M1 is sufficient to induce membrane deformation.**

**A**: Confocal LSM image of a typical GUV composed of DOPC:cholesterol:DOPS 50:20:30 molar ratios, in the presence of 10 µM M1C-Alexa488 (green channel). The lipid bilayer was labeled with 0.01 mol% Rhodamine-DOPE (red channel).

**B**: Confocal LSM image of a typical GUV with the same composition as in panel A, in the presence of 10 µM M1N-Alexa488 (green channel). The lipid bilayer was labeled with 0.01 mol% Rhodamine-DOPE (red channel).

All GUVs contained 150 mM sucrose in their lumen and were suspended in a phosphate buffered protein solution (pH 7.4) with similar osmolarity (see Materials and Methods).

Scale bars are 5 µm. Images were acquired at 23°C.

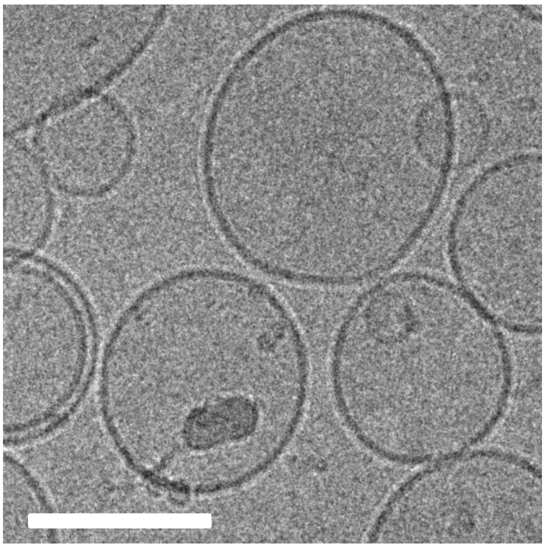

**FIGURE S3: Protein-free controls for membrane deformation.**

Typical cryo-TEM image of liposomes composed of 40 mol% DOPS in PBS pH 7.4, without M1. Few multilamellar vesicles can be observed but the shape of the liposomes is generally spherical. The scale bar is 100 nm.

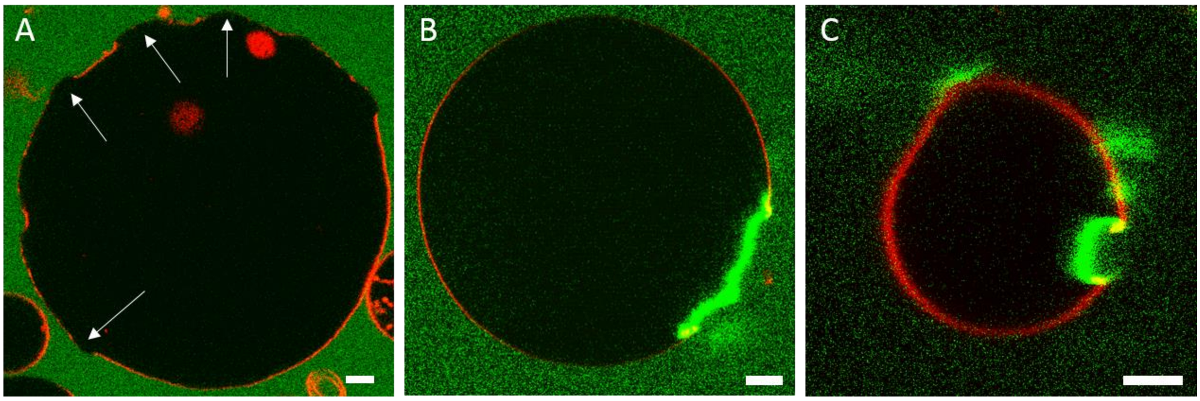


**FIGURE S4: Phase separated GUVs interacting with M1.**

**A**: Example of a GUV composed of cholesterol:DPPG:DSPC:DOPC 10:15:30:45 molar ratios (same as Fig. 3 in the main text), displaying outward budding of ordered domains (see arrows), before protein addition. Phase separation was observed by labeling the vesicles with Rhodamine-DOPE 0.01 mol% (red channel, strongly enriched in the disordered phase) and introducing water-soluble Alexa Fluor 488 succinimidyl ester in the outer milieu of the vesicles (green channel).

**B-C**: High contrast visualization of the images shown in Fig. 3 B and C, respectively. The green signal was enhanced offline so to evidence the presence of unbound M1-Alexa488 (green channel) in the outer milieu of the GUVs and its absence in the lumen.

Scale bars are 5 µm. Images were acquired at 23°C.

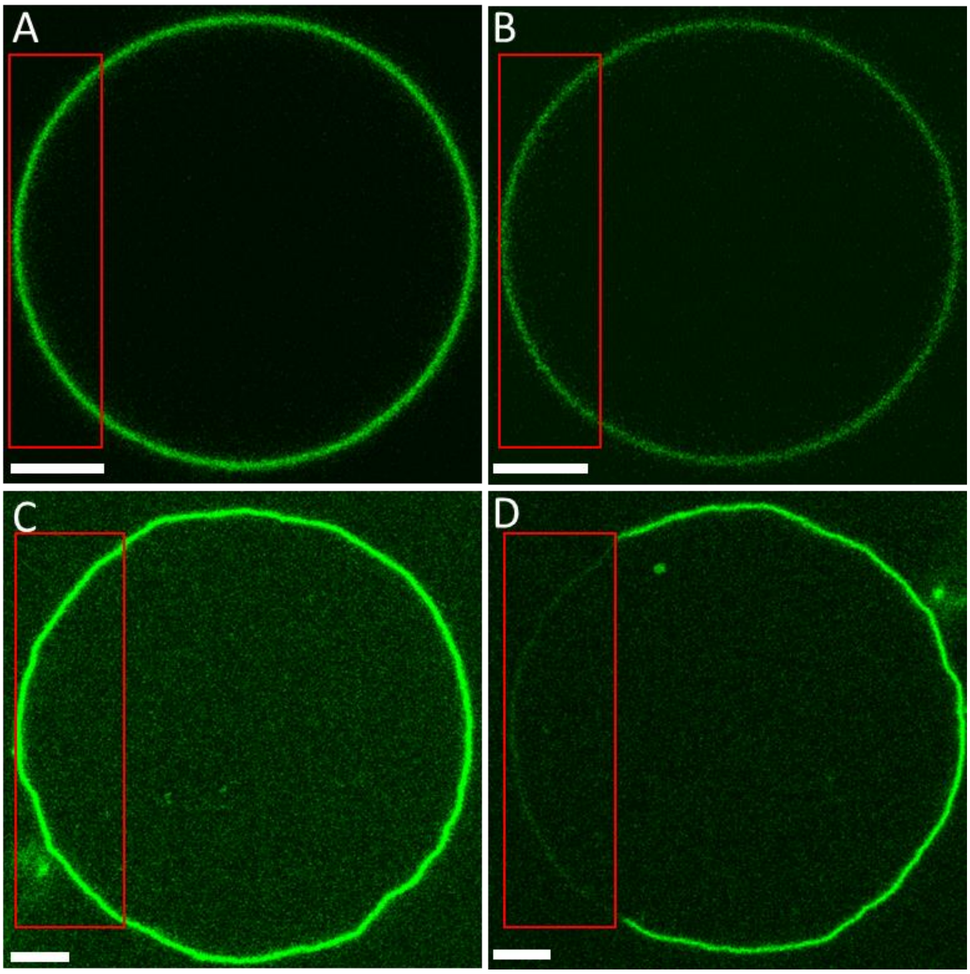


**FIGURE S5: Comparison of M1 and Annexin V binding. Annexin V is characterized by faster dynamics and does not induce significant membrane deformation**

**A-B**: Representative bleaching/recovery experiment for GUVs composed of DOPC:DOPS 70:30 molar ratio, in the presence of 20 µM FITC-Annexin V. It is worth noting that the affinity of Annexin V for phosphatidylserine-containing membrane (1, 2) is similar or even higher than that of M1 (3, 4). Panel A shows a typical vesicle before bleaching the fluorescent protein in the region enclosed in the red rectangle. The vast majority of the examined GUVs does not show visible membrane deformation, although these samples were observed in slightly hyperosmotic buffer conditions (i.e. 150 mM sucrose in the lumen of the GUVs; 180 mM glucose, 3 mM CaCl2, 20 mM NaCl in the external milieu). Panel B shows the same vesicle 5 s after a 10 s bleaching cycle at high laser power. Complete fluorescence signal recovery within few seconds suggests significantly fast protein lateral diffusion.

**C-D**: Bleaching experiment for GUVs with the same composition as in panels A-B, in the presence of 10 µM Alexa488-M1. Panel C shows a typical vesicle before bleaching the region enclosed in the red rectangle. Circa 50% of the examined GUVs displayed a deviation from spherical shape. In order to exclude that low membrane tension might play a role in membrane deformation, these GUVs were prepared in slight hypoosmotic conditions (i.e. 180 mM sucrose in the lumen of the GUVs and 2-fold diluted PBS buffer in the external milieu). Panel D shows the same vesicle 5 minutes after a 10 s bleaching cycle at high laser power. The absence of fluorescence signal recovery suggests the protein lateral diffusion is much slower compared to the case of Annexin V (Panels A-B). On the other hand, fast fluorescence signal recovery was observed for a fluorescent lipid probe (data not shown).

Imaging was performed at 25 °C. Scale bars are 5 µM.

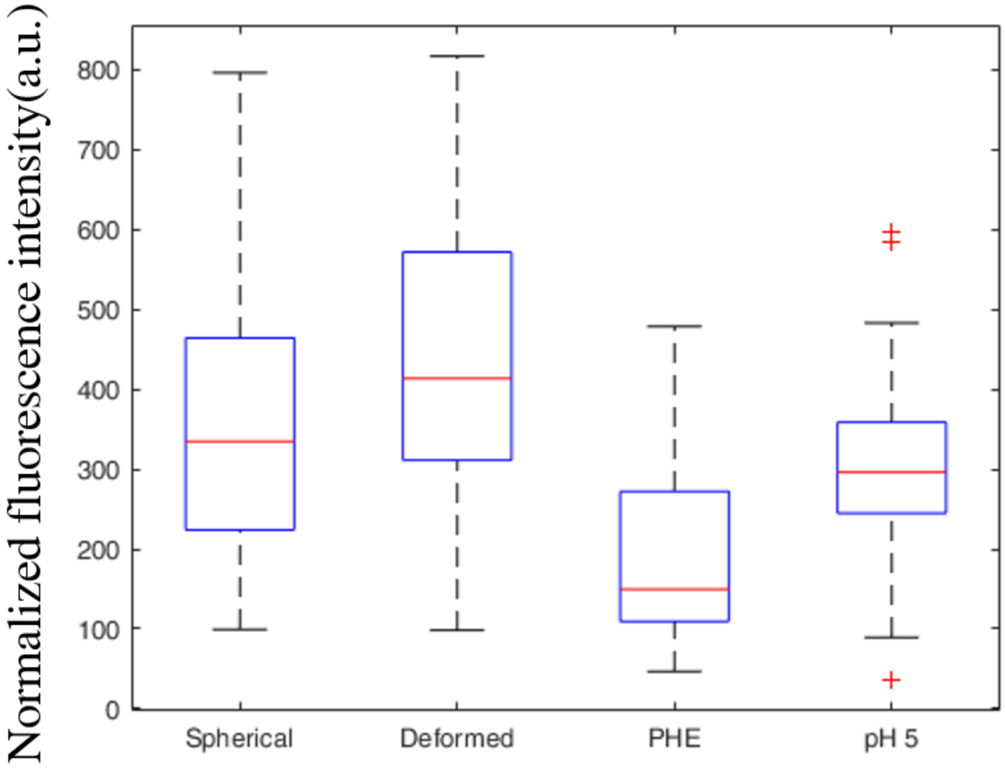


**FIGURE S6: sFCS analysis of M1 binding to spherical and deformed GUVs (see also Fig. 6).** GUVs composed of 30 mol% DOPS and 70 mol% DOPC were incubated for 30 minutes with 10 µM M1-Alexa488. The categories “Spherical” and “Deformed” refer to measurements in GUVs from samples prepared at pH 7.4, in the absence of PHE. In these conditions, ca. 50% of the GUVs are clearly non-spherical (see Table 1 of main text). The category “PHE” refers to spherical GUVs in samples prepared at pH 7.4, using 100 µM PHE. In these conditions, ca. 90% of the GUVs are clearly spherical (see Table 1). The category “pH 5” refers to spherical GUVs in samples prepared at pH 5, in the absence of PHE. In these conditions, ca. 90% of the GUVs are clearly spherical (see Table 1). Details of sample preparation are described in the Materials and Methods section. sFCS measurements were performed on 16-34 GUVs from two independent sample preparations and the results were pooled together. Each measurement provided M1-Alexa488 normalized brightness values and diffusion times (shown in Fig. 6 of the main text) and normalized fluorescence intensities (shown here). The category “Spherical” is significantly different from the category “PHE” (t-test $p<0.01$), but not from the other two categories (t-test $p>0.05$).

1. **Rosenbaum S, Kreft S, Etich J, Frie C, Stermann J, Grskovic I, Frey B, Mielenz D, Poschl E, Gaipl U, Paulsson M, Brachvogel B.** 2011. Identification of novel binding partners (annexins) for the cell death signal phosphatidylserine and definition of their recognition motif. J Biol Chem **286:**5708-5716.
2. **Kim S, Bae SM, Seo J, Cha K, Piao M, Kim SJ, Son HN, Park RW, Lee BH, Kim IS.** 2015. Advantages of the phosphatidylserine-recognizing peptide PSP1 for molecular imaging of tumor apoptosis compared with annexin V. PLoS One **10:**e0121171.
3. **Brevnov V, Fedorova N, Indenbom A.** 2016. Formation of the layer of influenza A virus M1 matrix protein on lipid membranes at pH 7.0. Russian Chemical Bulletin **65:**2737-2744.
4. **Hofer CT, Di Lella S, Dahmani I, Jungnick N, Bordag N, Bobone S, Huang Q, Keller S, Herrmann A, Chiantia S.** 2019. Structural determinants of the interaction between influenza A virus matrix protein M1 and lipid membranes. Biochim Biophys Acta Biomembr **1861:**1123-1134.
